# CRISPR/Cas9 screenings reveal the role of STX1A and CDK1 in Cathepsin G entering and killing colorectal cancer cells

**DOI:** 10.1101/2025.09.24.678097

**Authors:** Yuxiang Wang, Valery Rozen, Trang Dinh, He Li, Yamu Li, Zhenghe Wang

## Abstract

Neutrophils are the major populations of white blood cells and have been reported to facilitate cancer metastasis. Meanwhile, emerging evidence has recently suggested the anti-cancer role of neutrophils. Our previous study revealed that CB-839 and 5-FU-treated colorectal cancer (CRC) tumors recruited neutrophils and induced neutrophil extracellular traps (NETs). Cathepsin G (CTSG), which is released during NET formation, enters CRC cells through the receptor for advanced glycation end products (RAGE) and cleaves 14-3-3ε to promote apoptosis. However, the detailed mechanism underlying CTSG’s anti-tumor function remains less studied. In this study, we report that CTSG enters CRC cells through RAGE-mediated endocytosis. Knocking out RAGE or inhibiting endocytosis blocks CTSG from entering CRC cells and attenuates CTSG-induced apoptosis. Furthermore, the clathrin coat assembly complex and SNARE proteins were enriched in an arrayed CRISPR/Cas9 screening targeting human membrane trafficking genes. Knocking out SNARE protein STX1A prevents the spread of CTSG in CRC cells and the induction of cleaved PARP. A pooled genome-wide CRISPR/Cas9 screening further identifies the role of CDK1 in the NET-induced killing of CRC cells. Inhibiting CDK1 protected CRC cells from killing by CTSG. Our study reveals novel mechanisms by which CTSG enters and kills CRC cells.

## 1 Introduction

Neutrophils are fundamental for primary immune defense and account for 50-70% of white blood cells in human blood^1^. Neutrophils eliminate invading agents mainly through degranulation, phagocytosis, and neutrophil extracellular traps (NETs)^2^. However, eliminating invaded pathogens by neutrophils comes at a cost as the reactive oxygen species (ROS), neutrophil granules, cytokines, and NETs generated and released during the progress result in neutrophil-driven toxicity^3^. NETs can cause tissue damage in inflammatory diseases like anti-neutrophil cytoplasmic antibody-associated autoimmune vasculitis, systemic lupus erythematosus, and rheumatoid arthritis^4^. In addition to inflammatory diseases, neutrophils are involved in the tumor immune microenvironment. Many studies have reported the role of NETs in promoting tumor metastasis and mediating drug resistance^5-11^.

A growing body of evidence also supports the anti-tumor role of neutrophils recently^12-14^. We recently demonstrated that chemotherapies induce NETs and augment their anti-tumor effect^15^. We found that *PIK3CA*, which encodes the p110α catalytic subunit of phosphatidylinositol-4,5-bisphosphate 3-kinase (PI3K), renders colorectal cancers (CRCs) dependent on glutamine^16;17^. We further showed that a combination of glutaminase inhibitor CB-839 and 5-FU induces the regression of *PIK3CA*-mutant CRCs^18^. Mechanistically, the drug combination induced the expression of IL-8 in *PIK3CA* mutant CRC cells, leading to the recruitment of neutrophils in tumors^15^. Additionally, the drug combination increased the levels of ROS in neutrophils, thereby inducing NETs^15^. NET-associated Cathepsin G (CTSG) then entered CRC cells through the receptor for advanced glycation end products (RAGE) on the cell membrane, leading to the cleavage of 14-3-3ε proteins, releasing Bcl-2-associated X protein (Bax) to mitochondria, and triggering apoptosis in CRC cells^15^.

However, the exact mechanisms underlying how CTSG enters and kills CRC cells remain poorly understood. In this study, we showed that CTSG entered CRC cells through RAGE-mediated endocytosis. Knocking out RAGE or inhibiting endocytosis blocked CTSG from entering CRC cells and attenuated CTSG-induced apoptosis. Furthermore, we applied arrayed and pooled CRISPR/Cas9-based screenings to identify potential regulators mediating cell entry and cell killing of CTSG. The clathrin coat assembly complex and SNARE proteins were enriched in arrayed CRISPR/Cas9 screening targeting membrane trafficking genes. Knocking out SNARE protein STX1A prevented the spread of CTSG in CRC cells and the induction of cleaved PARP. Moreover, pooled genome-wide CRISPR/Cas9 screening indicated that KO of CDK1 relieved NET-induced apoptosis in CRC cells. CDK1 inhibitor Ro-3306 protected CRC cells from CTSG-induced apoptosis. Our study elucidates novel mechanisms by which CTSG enters and kills CRC cells.

## 2 Materials and methods

### Cell lines and cell culture

HCT116 (#CCL-247), DLD1 (#CCL-221), and 293T (#CRL-3216) cells were purchased from ATCC. HCT116 cells were cultured with McCoy’s 5a medium with 10 % fetal bovine serum (FBS). DLD1 cells were cultured with RPMI-1640 medium with 10 % FBS. 293T cells were cultured with DMEM medium with 10 % FBS. All cells were incubated under 5% CO_2_ at 37 °C. RAGE-KO HCT116 and DLD1 cells were generated previously^15^. STX1A-KO HCT116 and DLD1 cells were generated by CRISPR/Cas9 system as described previously^19^.

### Compounds

Dynasore (#S8047) and Ro-3306 (#S7747) were purchased from Selleckchem. All powders were dissolved in dimethyl sulfoxide (DMSO) to make a 10 mM stock solution. The 10 mM stock solution was aliquoted and stored at -20 °C.

### Caspase-3/7 activity assay

Caspase-Glo 3/7 assay system was purchased from Promega (#G8091). The assay was conducted referring to the standard protocol for cells cultured in a 96-well plate from Promega.

Briefly, plates containing cells and Caspase-Glo 3/7 reagent were equilibrated to room temperature before use. 100 μL of Caspase-Glo 3/7 reagent was added into wells containing 100 μL of medium. The plate was gently shaken at 300-500 rpm for 30 seconds, incubated at room temperature for 1 hour, and the luminescence of each sample and negative control wells was measured.

### Co-immunoprecipitation assay

His tag fused RAGE was overexpressed in 293T cells. Cells were lysed with the lysis buffer (50 mM NaH_2_PO_4_, 300 mM NaCl, 10 mM imidazole, pH 8.0) and incubated with 2 μg CTSG at 4 °C for 2 hours. His-tag fused RAGE was then enriched by Ni-NTA beads, and CTSG was co-immunoprecipitated. The Ni-NTA agarose beads were pelleted by spin down and washed 3 times. Immuno-complexes were eluted with SDS loading buffer by boiling for 10 min and subjected to Western blotting.

### Western blotting

Cultured cells were lysed with lysis buffer (50 mM Tris pH 7.4, 150 mM NaCl, 5 mM EDTA, 0.5% NP40, 0.1% SDS, 1 mM PMSF, complete Protease Inhibitor Cocktail, and phosphatase inhibitors). Then, lysates were cleared by centrifugation (14,000 rpm, 10 min), and protein concentration was determined by the BCA protein assay kit. Equal amounts of total protein were used for immunoblotting.

### Immunofluorescent staining

HCT116 and DLD1 cells were seeded on glass slides in a 24-well plate. Dynasore and CTSG were then added to the medium for the indicated time. The treated cells were fixed with 4% paraformaldehyde. The slides were then blocked with 0.1% Triton-X100 in 5% BSA for 30 min at room temperature, followed by incubation with a primary antibody overnight. After the slides were washed 3 times by PBST, a secondary antibody prepared in blocking buffer was added and incubated for 1 hour at room temperature. After washing 3 times with PBST, slides were mounted with a prolonged anti-fade solution with DAPI and stored at -20 °C before visualization.

### Arrayed CRISPR/Cas9 screening

The LentiArray™ Human Membrane Trafficking CRISPR Library (#A42272, Thermo Fisher) was employed to induce KO membrane trafficking-related genes in HCT116 cells stably expressing Cas9. HCT116-Cas9 cells were seeded in 12-well plates at 150000 cells/well. 50 μL of arrayed lentivirus targeting 141 membrane trafficking-related genes with RAGE targeting sgRNAs as positive control and scramble sgRNAs as negative control the next day. Virus-infected cells were reseeded in 24-well plates at 150000 cells/well without FBS after puromycin selection for 3 days, and 5 ng/μL of CTSG was added. Cells were lysed with SDS loading buffer, boiled for 10 min, and subjected to Western blotting. The level of cleaved PARP was evaluated as a marker of cell entry of CTSG.

### Pooled CRISPR/Cas9 screening

The genome-wide CRISPR screening was performed as described previously^20^. Briefly, the Human CRISPR Knockout Pooled Library (Brunello, # 73178-LV, Addgene) was employed to induce genome-wide knockout in DLD1 cells stably expressing Cas9. DLD1-Cas9 cells were transduced with the lentiviral-pooled sgRNA library at a low multiplicity of infection (MOI ≈0.3). Infected cells were cultured with puromycin for 3 days and then treated with condition medium from neutrophils or DLD1 cells as a negative control. After treatment for 1 day, genomic DNA was extracted with a QIAamp DNA Blood Maxi Kit (QIAGEN). The sgRNA fragments were amplified by a two-step polymerase chain reaction (PCR) with the indicated primers as below. The PCR products were sequenced with the Illumina HiSeq X ten System. Sequencing reads were aligned to the sgRNA library sequences by Bowtie 2 and analyzed with MAGeCK. The counts of sgRNA reads were normalized, and differential analysis among the samples was performed.

Step 1 Fwd, CTTTCCCTACACGACGCTCTTCCGATCTTTGTGGAAAGGACGAAACACCG. Step 1 Rev, GACTGGAGTTCAGACGTGTGCTCTTCCGATCTTCTACTATTCTTTCCCCTGCACTGT. Step 2 Fwd, AATGATACGGCGACCACCGAGATCTACACTATAGCCTACACTCTTTCCCTACACGACGCTCTTCC. Step 2 Rev, CAAGCAGAAGACGGCATACGAGATCGAGTAATGTGACTGGAGTTCAGACGTGTGCTCTTC.

### Statistics

GraphPad Prism software was used to create the graphs. Data are plotted as mean ± SD. For 2 group comparison, we applied the 2-tailed t test to compare the means between 2 groups, assuming unequal variances. P < 0.05 is defined as statistically significant.

## 3 Results

### 3.1 CTSG enters cancer cells through endocytosis

Our previous study showed that CTSG entered CRC cells in a RAGE-dependent manner^15^. However, the detailed mechanism underlying RAGE-mediated cell entry of CTSG remains largely unknown. It has been reported that RAGE-mediated cell entry of high mobility group box 1 protein (HMGB1)^21-23^, Aβ^24^, and endotoxin (lipopolysaccharide, LPS)^23;25^ through endocytosis. To test whether CTSG also enters CRC cells through endocytosis, we incubated CRC cell HCT116 with CTSG and endocytosis inhibitor Dynasore for 2 hours. CTSG entered HCT116 cells and spread evenly in HCT116 cells (Fig. 1A). However, CTSG was localized around the cell membrane when Dynasore inhibited endocytosis (Fig. 1A). The data suggest that blocking endocytosis attenuated the spread of CTSG within the cells. As CTSG induced apoptosis after entering cancer cells^15^, we interrogated apoptosis signaling in HCT116 cells. CTSG increased the activity of caspase-3/7 (Fig. 1B) and the levels of cleaved PARP (Fig. 1C) in HCT116 cells, whereas Dynasore prevented CTSG-induced caspase-3/7 activity and cleaved PARP levels (Fig. 1B and 1C). To further verify that CTSG enters cancer cells through endocytosis, we stained early endosome marker, early endosome antigen 1 (EEA1), together with CTSG. CTSG attached to cell membranes at 10 minutes after incubation and co-localized with EEA1 at 20 and 30 minutes after incubation (Fig. 1D). CTSG started to dissociate with EEA1 at 40 minutes after incubation (Fig. 1D). These results suggest that endocytosis is essential for the spreading of CTSG within cells.

**Figure 1.**
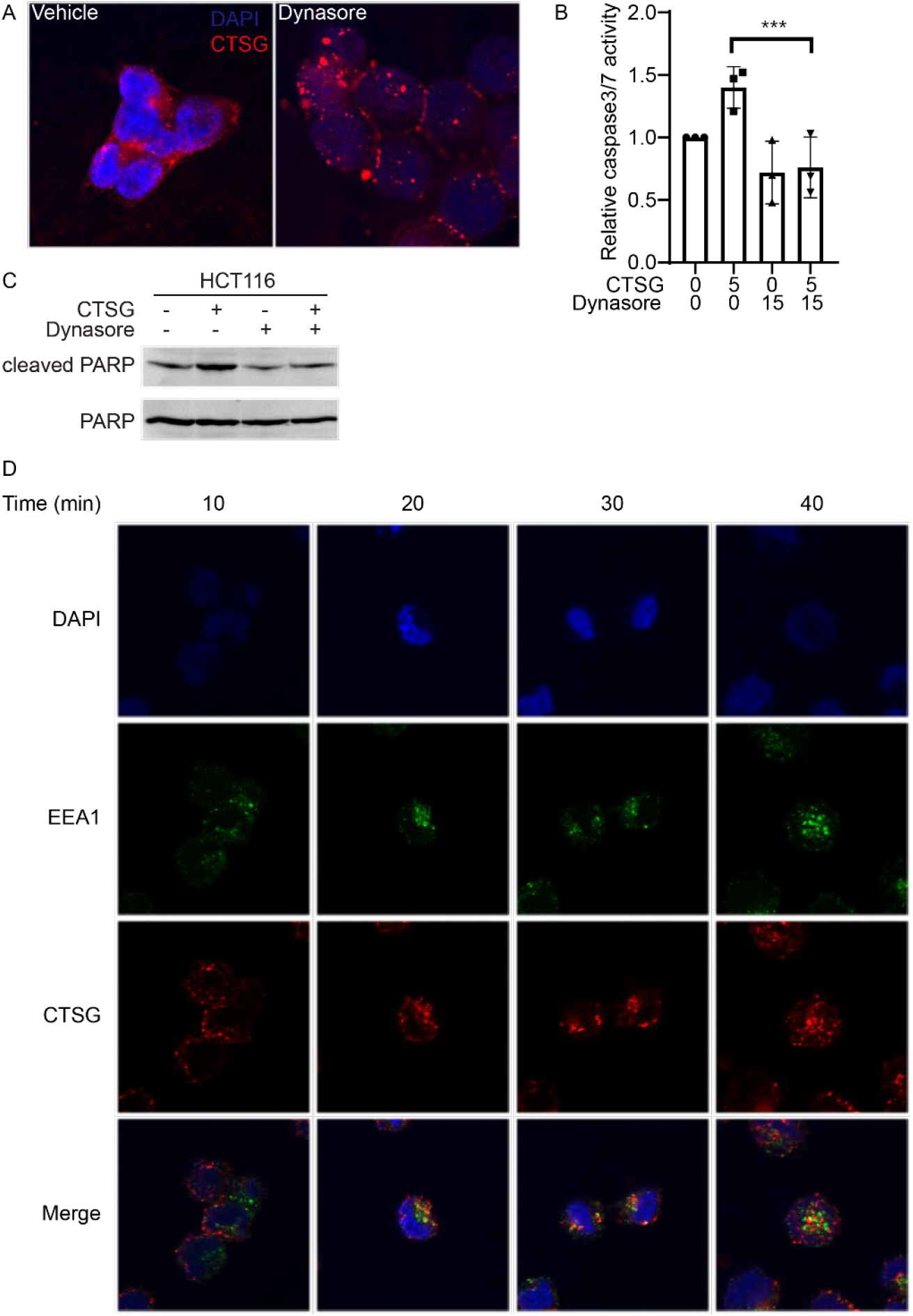
CTSG enters cancer cells through endocytosis. (A) HCT116 cells were treated with 5 ng/μL of CTSG for 2 hours with or without 15 μM Dynasore. The data presented are representative images of immunofluorescent staining of an anti-CTSG antibody. (B-C) HCT116 cells were treated with 5 ng/μL of CTSG for 16 hours with or without 15 μM Dynasore. Relative caspase-3/7 activity is shown in B (n = 3), and Western blots of cleaved PARP are shown in C. (D) HCT116 cells were treated with 5 ng/μL of CTSG for the indicated time. The data presented are representative images of immunofluorescent staining of anti-CTSG and anti-EEA1 antibodies.

### 3.2 CTSG enters cancer cells in a RAGE-dependent manner

To study whether RAGE is required for CTSG entering cancer cells, we incubated RAGE-KO and parental HCT116 cells with CTSG. CTSG entered parental HCT116 cells but not RAGE-KO HCT116 cells at 4 hours after incubation (Fig. 2A). Accordingly, CTSG decreased cell viability in HCT116 cells rather than RAGE-KO HCT116 cells (Fig. 2B). As RAGE is a nucleic acid receptor and CTSG binds to decondensed DNA released during NETs formation^26^, we further tested whether CTSG directly interacts with RAGE. We overexpressed recombinant RAGE protein fused with a His tag in HEK293T cells and incubated the cell lysates with CTSG. CTSG and RAGE were precipitated by Ni-NTA agarose beads. Although CTSG is bound to the empty beads slightly, His-tagged RAGE-bound beads precipitated more CTSG in the system (Fig. 2C). These results suggest that CTSG interacts with RAGE and mediates the cell entry of CTSG.

**Figure 2.**
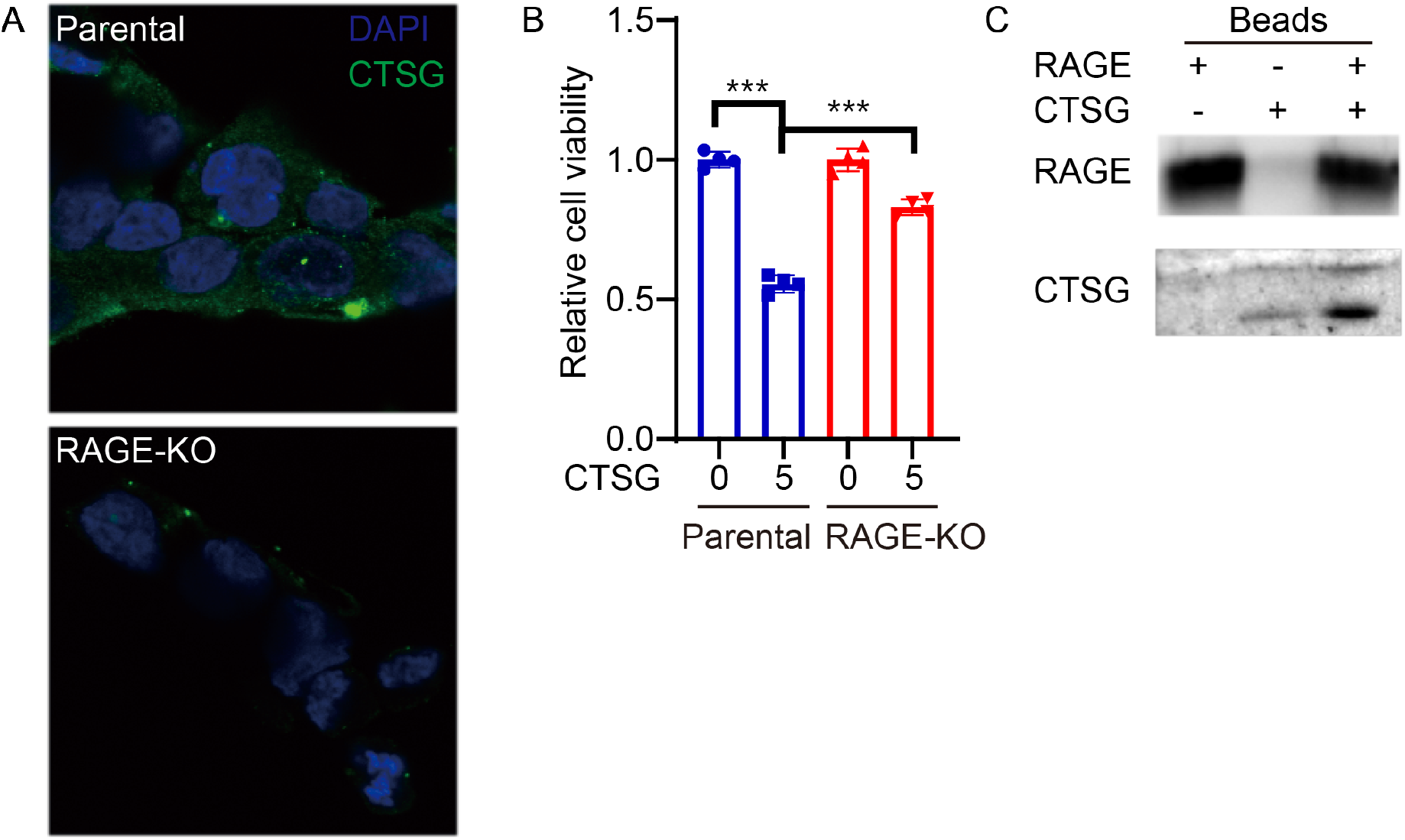
CTSG enters cancer cells in a RAGE-dependent manner. (A) Parental and RAGE-KO HCT116 cells were treated with 5 ng/μL of CTSG for 4 hours. The data presented are representative images of immunofluorescent staining of an anti-CTSG antibody. (B) Parental and RAGE-KO HCT116 cells were treated with 5 ng/μL of CTSG for 16 hours. The data presented are relative cell viability (n = 3). (C) Plasmid expressing His-tag fused RAGE was transduced into HEK293T cells. HEK293T cells were lysed 48 hours after transduction, and CTSG was added to the cell lysis. RAGE and CTSG were precipitated with Ni-NTA beads and subjected to Western blot assay with anti-His tag and anti-CTSG antibody.

### 3.3 Arrayed CRISPR/Cas9 screening revealed that the vesicle organization pathway was important for cell entry of CTSG

CRISPR/Cas9-based screening has emerged as a powerful tool to attribute functional phenotypes to various gene perturbations^27^. To figure out which protein contributed to cell entry of CTSG through RAGE-mediated endocytosis, we conducted an arrayed CRISPR screening with a sgRNA library targeting 141 human membrane trafficking genes (Supplementary Table 1). The sgRNAs targeting RAGE were used as positive controls, and sgRNAs with scramble sequence as a negative control (NC). We set NC sgRNA as the baseline to show how CTSG increased cleaved PARP and set sgRNA targeting RAGE as the threshold as we had proved that knockout of RAGE prevented CTSG from entering cells. Basically, we seeded HCT116 in 12-well plates and infected these cells with virus-containing sgRNA targeting 141 human membrane trafficking genes, RAGE, and NC. After puromycin selection for 3 days, we reseeded the cells in 24-well plates with the FBS-free medium and treated the cells with CTSG overnight. The next day, we harvested the cell lysis for Western blotting using the level of cleaved PARP/Actin to indicate cell entry of CTSG. Compared to knocking out RAGE, knocking out 40 different genes reduced the CTSG-induced cleaved PARP to a greater level (Fig. 3A). Pathway and process enrichment analysis revealed that the 40 genes were enriched in pathways related to vesicle organization, regulation of calcium ion-dependent exocytosis, melanosome assembly, and membrane trafficking (Fig. 3B and Supplementary Table 2). To further understand the relationships between the enriched pathways, a network plot analysis was performed on the 40 genes and showed that they belonged to pathways that interacted mainly with each other (Fig. 3C).

**Figure 3.**
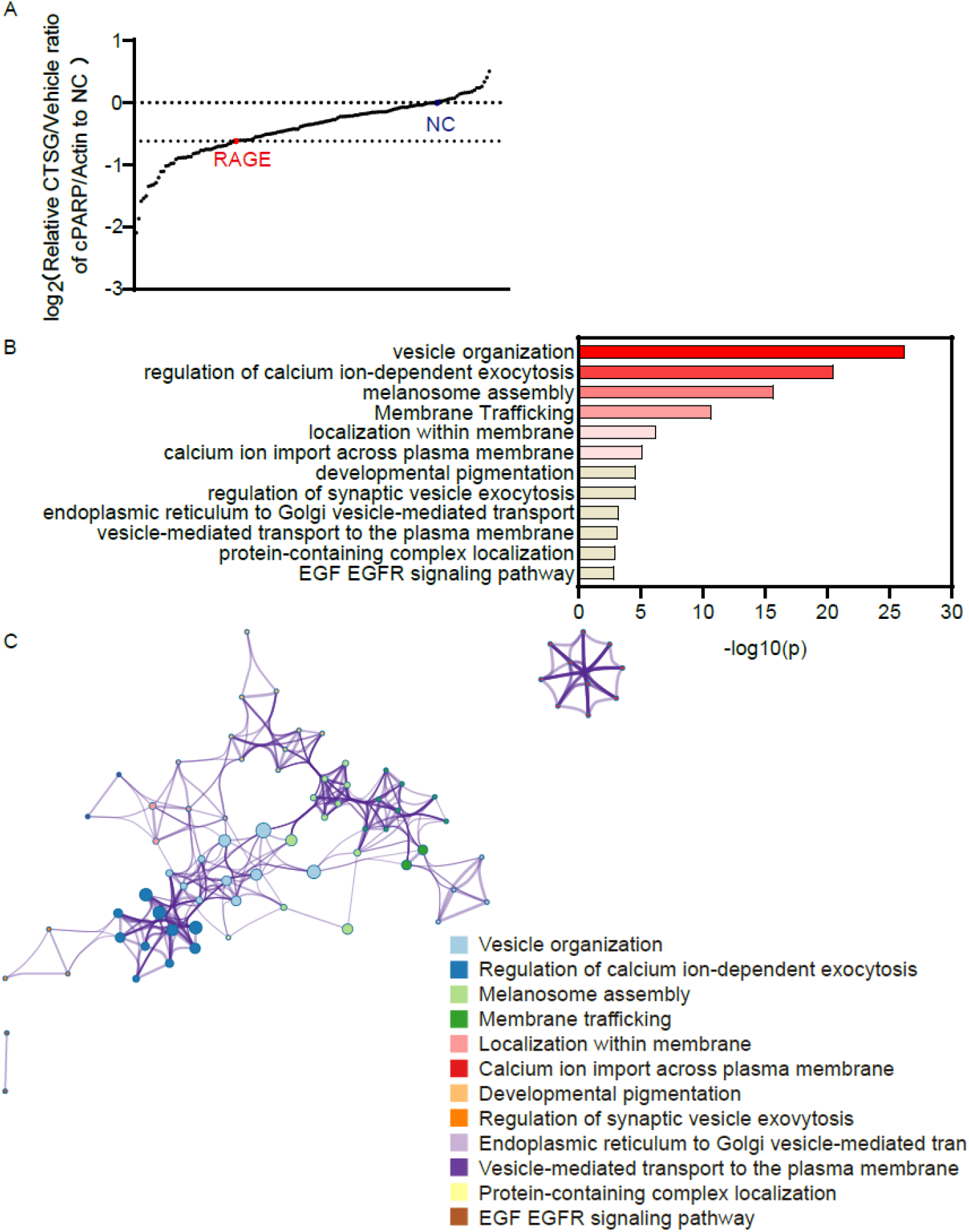
Arrayed CRISPR/Cas9 screening revealed that the vesicle organization pathway was important for cell entry of CTSG. (A) Cas9 expressing HCT116 cells were infected with 50 μL of arrayed gRNA lentivirus targeting 141 human membrane trafficking genes, RAGE, and negative control (NC). Infected cells were selected with puromycin for 3 days. The cells were then reseeded and treated with 5 ng/μL of CTSG for 16 hours. Cleaved PARP and Actin were detected with Western blot assay. Cleaved PARP levels were normalized with Actin, and the relative ratio of cleaved PARP in CTSG-treated wells against vehicle control wells to NC was calculated and plotted. (B-C) The enrichment of the top 40 genes was analyzed using Metascape (http://metascape.org/)^34^. The pathway enrichment list is shown in B, and the relationships of the enriched pathways are shown in C.

### 3.4 STX1A-mediated vesicle fusion is required for CTSG-induced apoptosis

To figure out the mechanism by which CTSG enters cancer cells, we studied the protein-protein interactions of the 40 genes. The protein-protein interactions were mainly enriched among the clathrin coat assembly complex and SNARE proteins (Fig. 4A). The clathrin coat assembly complex is involved in endocytosis and Golgi processing. Dynamin inhibitor Dynasore blocked CTSG from entering cells, further supporting the conclusion that endocytosis is essential for cell entry of CTSG. The SNARE proteins are necessary for protein trafficking and fusion of intracellular transport vesicles^28^. We then aimed to verify whether SNARE proteins affect CTSG entry into cells and spread within cells. As STX1A is found at endosomes^29^ and is among the top 10 enriched genes, we thus knocked out the STX1A gene in HCT116 and DLD1 cells and tested cleaved PARP levels induced by CTSG and CTSG distribution in cells. Knocking out STX1A attenuated CTSG-induced cleaved PARP in both HCT116 and DLD1 cells (Fig. 4B & 4C). While CTSG spread evenly in parental DLD1 cells, CTSG accumulated more in STX1A-KO DLD1 cells, indicating that knocking out STX1A blocked the traffic of CTSG in the endosome (Fig. 4D).

**Figure 4.**
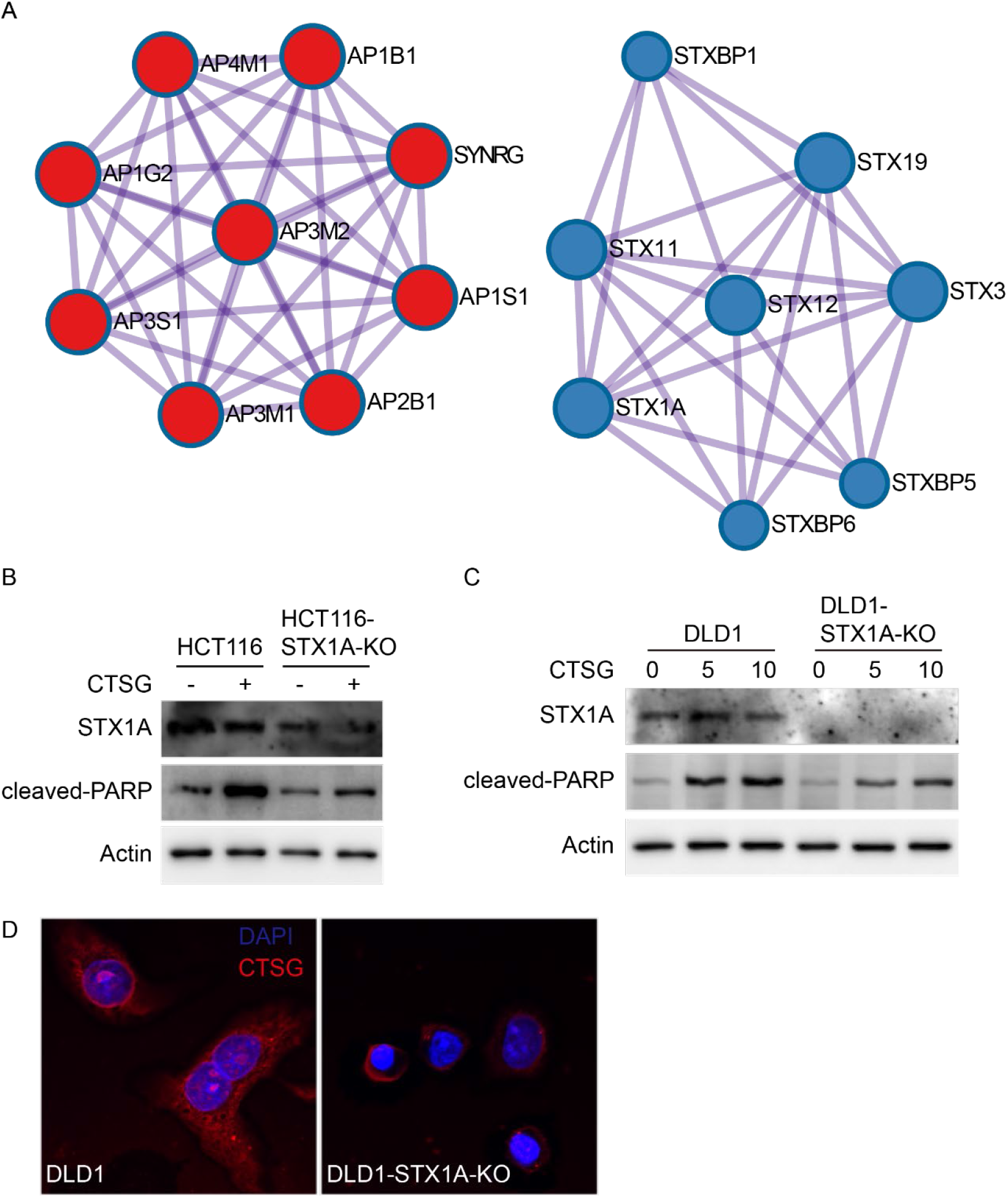
STX1A-mediated vesicle fusion is required for CTSG-induced apoptosis. (A) The protein-protein interaction network of the top 40 genes was analyzed using Metascape (http://metascape.org/). The data presented are two main protein-protein interaction networks. (B-C) Parental and STX1A-KO HCT116 and DLD1 cells were treated with CTSG at the indicated concentrations for 16 hours. Data presented are Western blots of STX1A, cleaved PARP, and Actin in HCT116 cells (B) and in DLD1 cells (C). (D) Parental and STX1A-KO DLD1 cells were treated with 5 ng/μL of CTSG for 4 hours. The data presented are representative images of immunofluorescent staining of an anti-CTSG antibody.

### 3.5 Pooled genome-wide CRISPR/Cas9 screening showed nucleus export pathways and nuclear transport pathways were enriched by GO analysis

To reveal the details of how CTSG released during NETs formation kills cancer cells, we conducted a genome-wide CRISPR/Cas9 screening in colorectal cancer cell line DLD1 treated with neutrophil conditional medium containing CTSG. By sequencing DLD1 cells treated with neutrophil conditional medium and DLD1 conditional medium as a control, we analyzed genes significantly enriched or decreased in DLD1 cells treated with neutrophil conditional medium. GO analysis showed that multiple nucleus export pathways and nuclear transport pathways are involved in CTSG-mediated cancer cell-killing function (Fig. 5A).

**Figure 5.**
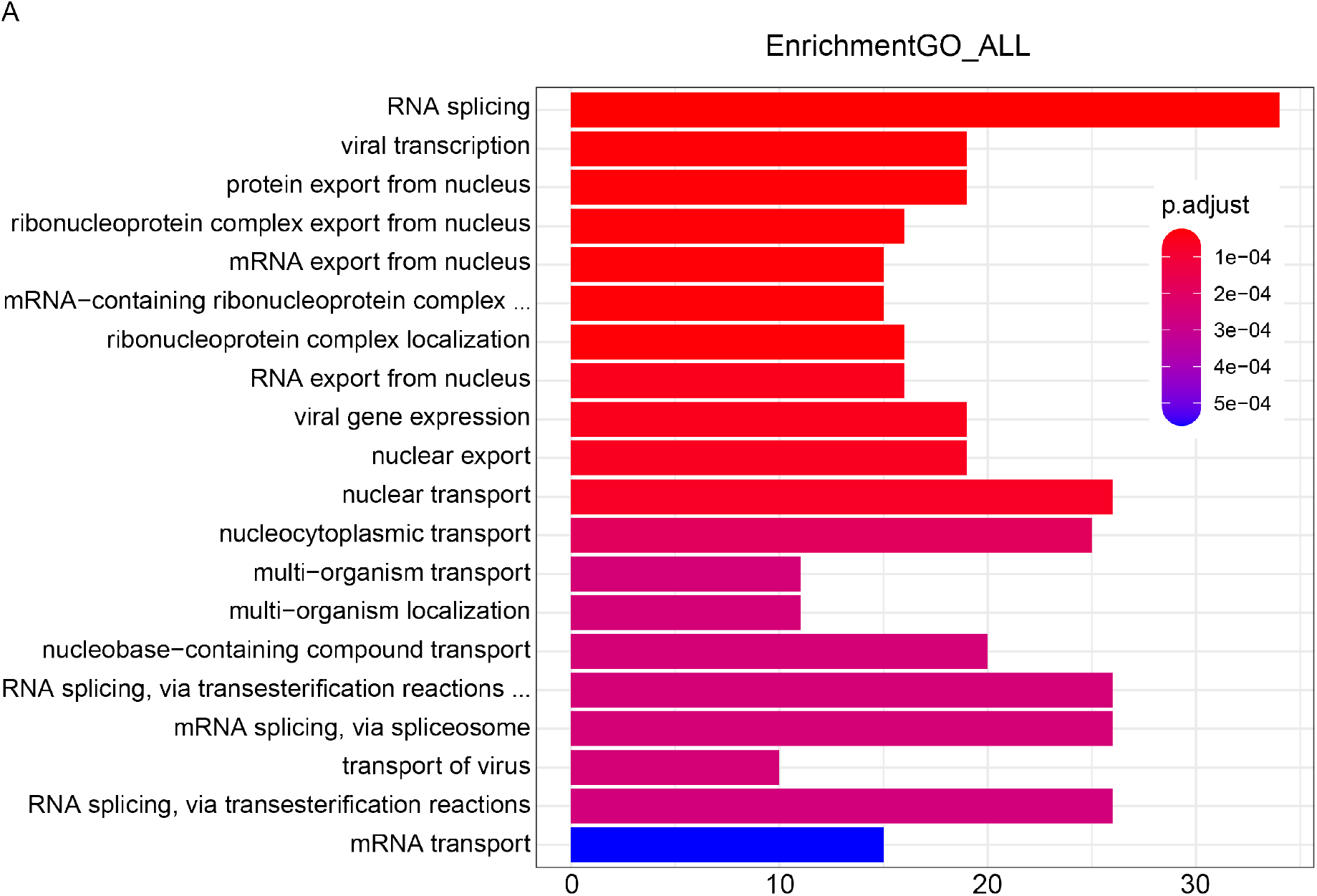
Pooled genome-wide CRISPR/Cas9 screening showed nucleus export pathways and nuclear transport pathways were enriched by GO analysis. (A) Cas9-expressing DLD1 cells were infected with genome-wide pooled gRNA lentivirus. Infected cells were selected with puromycin for 3 days. The cells were then reseeded and treated with NETs condition medium or DLD1 condition medium for 16 hours. Genomic DNA was extracted from DLD1 cells after treatment, and gRNA sequences were amplified by PCR. PCR products were purified and subjected to next-generation sequencing. The enriched genes were subjected to GO analysis. The data presented are the enriched pathways

### 3.6 CDK1 inhibitor attenuates the cell-killing function of CTSG

Interestingly, among the top 10 genes exhibiting significant changes, NDC80, CDK1, and INCENP are involved in cell cycle and closely linked to microtubule activity (Fig. 6A and supplementary Table 3). Notably, microtubules also serve as tracks for intracellular vesicle transport^30^. It has been reported that CDK1 phosphorylates epsin 1 and impairs its interaction with clathrin and other endocytic proteins, regulating the dynamics of clathrin-coated vesicles^31^. Given that CDK1 inhibitor, Ro-3306, is commercially available, we further studied the role of CDK1 in NETs/CTSG induced apoptosis. As shown in Fig. 6B, CDK1 inhibitor Ro-3306 protected HCT116 and DLD1 cells from CTSG-induced apoptosis in a dose-dependent manner. Accordingly, Ro-3306 attenuated CTSG-induced cleaved PARP (Fig. 6C) in a dose-dependent manner. As CDK1 also plays a key role in cell cycle progression through the G2/M phase transition^32^, we treated CRC cells with CTSG and nocodazole, which arrested cells in the G2/M phase. Nocodazole did not impact the cell-killing function of CTSG, indicating a cell cycle-independent role of CDK1 in mediating CTSG’s cell-killing function (Fig. S1A). However, Ro-3306 alone induced apoptosis when incubated with HCT116 and DLD1 cells for 72 hours, probably due to prolonged cell cycle arrest (Fig. S1B).

**Figure 6.**
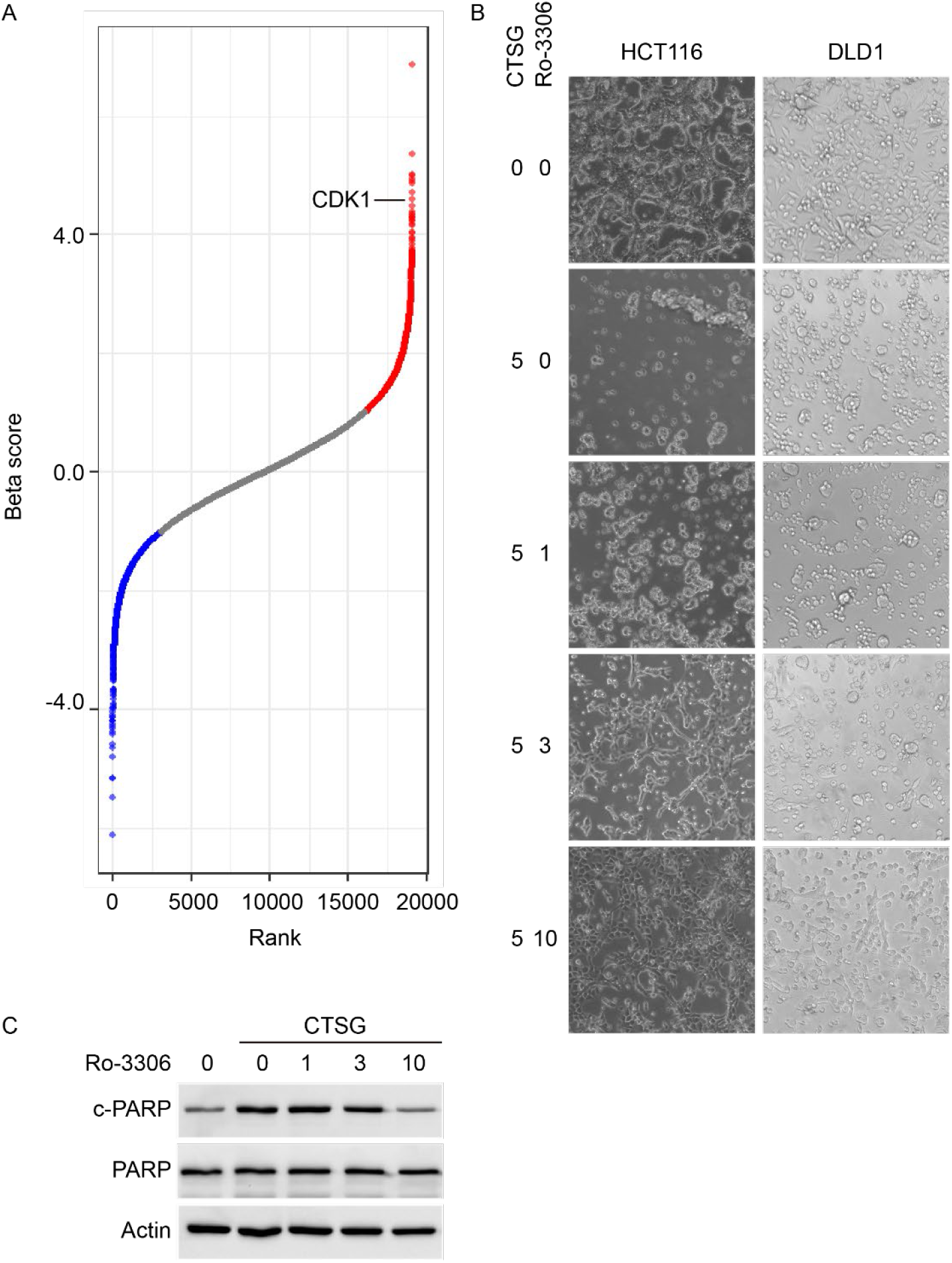
CDK1 inhibitor attenuates the cell-killing function of CTSG. (A) Cas9-expressing DLD1 cells were infected with genome-wide pooled gRNA lentivirus. Infected cells were selected with puromycin for 3 days. The cells were then reseeded and treated with NETs condition medium or DLD1 condition medium for 16 hours. Genomic DNA was extracted from DLD1 cells after treatment, and gRNA sequences were amplified by PCR. PCR products were purified and subjected to next-generation sequencing. The differentially expressed genes were plotted based on rank and Beta score. (B) HCT116 and DLD1 cells were treated with 5 ng/μL of CTSG for 24 hours with Ro-3306 at the indicated concentrations. The data presented are representative images of cellular morphology. (C) HCT116 cells were treated with 5 ng/μL of CTSG for 16 hours with Ro-3306 at the indicated concentrations (ng/μL). The data presented are Western blots of cleaved PARP.

## 4 Discussion

We revealed here that CTSG released by NETs enters cancer cells through RAGE-mediated endocytosis. Our studies demonstrated that RAGE and dynamin-dependent endocytosis are required for CTSG’s cell entry. Furthermore, arrayed CRISPR/Cas9 screening targeting membrane trafficking genes indicated that vesicle organization and transport pathways were involved in CTSG entering cancer cells. Specifically, the clathrin coat assembly complex and SNARE proteins were among the top enriched genes essential for CTSG entering cancer cells. Inhibiting clathrin and dynamin-dependent endocytosis by Dynasore and KO of SNARE protein STX1A blocked CTSG entering cells, thus preventing the induction of cleaved PARP by CTSG. It is reported that RAGE-mediated cell entry of HMGB1^21-23^, Aβ^24^, and endotoxin (lipopolysaccharide, LPS)^23;25^ through endocytosis, which is in accordance with our findings.

Neutrophils are involved in autoimmune diseases and chronic inflammatory diseases and are the main factor to cause tissue damage. Regulating CTSG’s cell-killing function may benefit both cancer patients and inflammatory disease patients. CDK1 stood out as a candidate target to relieve CTSG’s tumor-killing function in the pooled genome-wide CRISPR/Cas9 screening. CDK1 has been reported to regulate apoptosis through phosphorylating Bcl-xL/Bcl-2 and disabling their antiapoptotic activity^33^. CDK1 inhibitor Ro-3306 protected HCT116 and DLD1 cells from killing by CTSG when incubated for 24 hours. However, Ro-3306 itself induced apoptosis when incubated with HCT116 and DLD1 cells for 72 hours, probably due to prolonged cell cycle arrest. As nocodazole failed to protect CRC cells from being killed by CTSG, the cell cycle-independent function of CDK1 is involved. Determining the exact substrates of CDK1 involved in this process may help to develop therapies for autoimmune diseases and chronic inflammatory diseases caused by neutrophils.

We demonstrated that CTSG entered cancer cells through RAGE-mediated endocytosis, and CTSG co-localized with early endosomes in cancer cells. However, how CTSG is released into the cytoplasm from endosomes remains unclear. It is reported that macrophages and endothelial cells take up HMGB1-LPS complexes via RAGE-mediated endocytosis, and HMGB1 destabilizes lysosomes, releasing LPS to cytosolic caspase-11^23^. Further studies are needed to determine whether CTSG can destabilize endosomes or lysosomes.

---

**Supplementary Figure 1.**
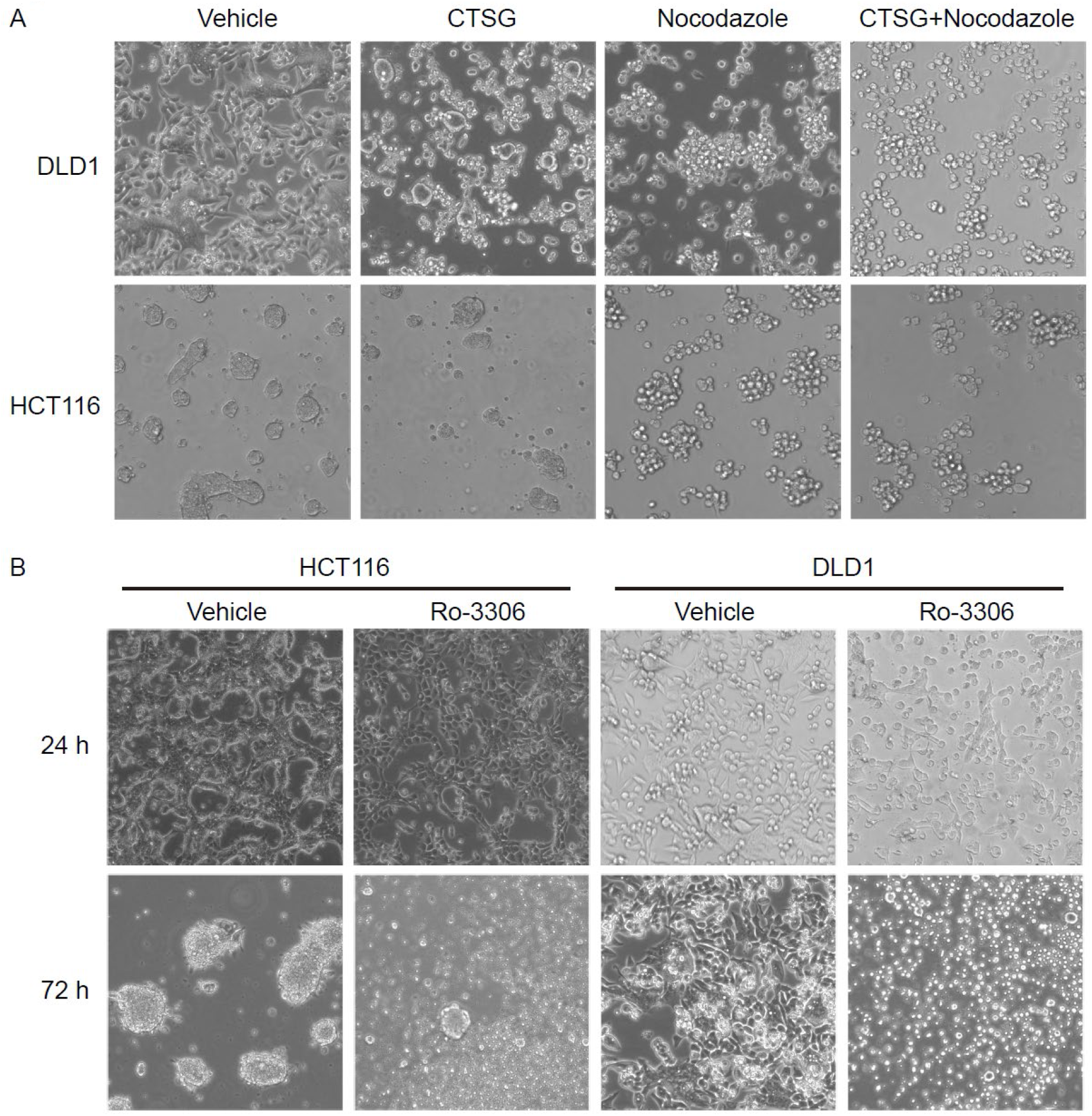
CDK1 inhibitor attenuates the killing function of CTSG in a cell cycle-independent manner. (A) HCT116 and DLD1 cells were treated with 5 ng/μL of CTSG, 1 ng/μL of nocodazole, or their combination for 24 hours. The data presented are representative images of cellular morphology. (B) HCT116 and DLD1 cells were treated with vehicle or 10 μM Ro-3306 for 24 or 72 hours. The data presented are representative images of cellular morphology.

**Supplementary table 1.**
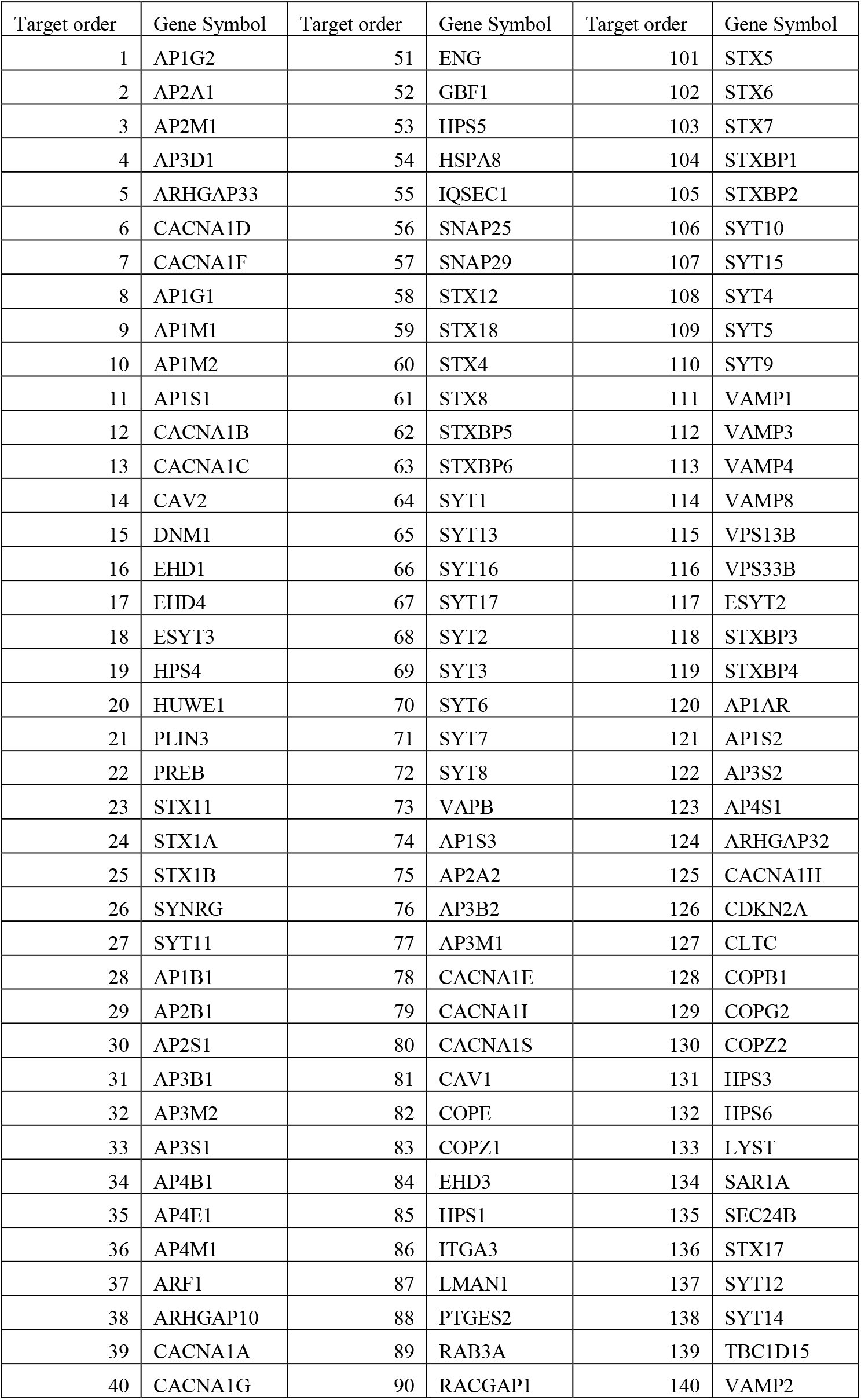
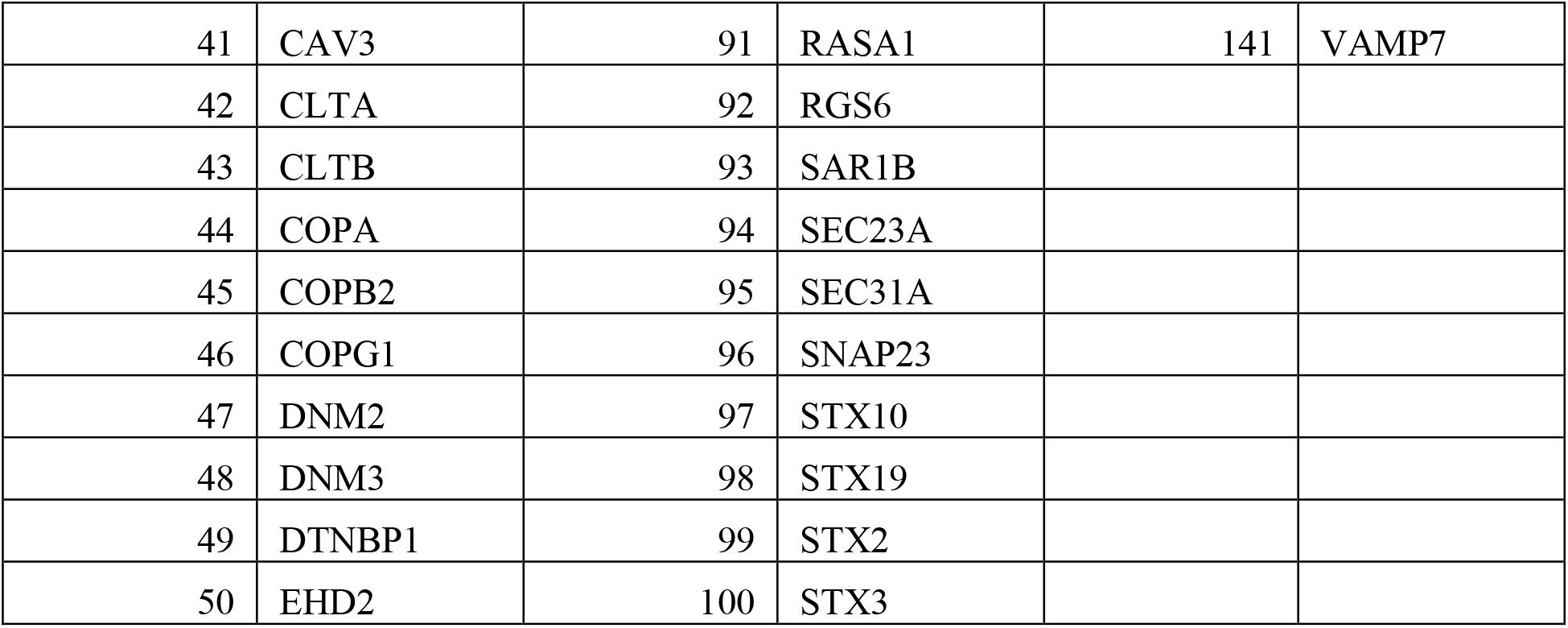
141 human membrane trafficking genes are included in the arrayed sgRNA library.

| Target order | Gene Symbol | Target order | Gene Symbol | Target order | Gene Symbol |
| --- | --- | --- | --- | --- | --- |
| 1 | AP1G2 | 51 | ENG | 101 | STX5 |
| 2 | AP2A1 | 52 | GBF1 | 102 | STX6 |
| 3 | AP2M1 | 53 | HPS5 | 103 | STX7 |
| 4 | AP3D1 | 54 | HSPA8 | 104 | STXBP1 |
| 5 | ARHGAP33 | 55 | IQSEC1 | 105 | STXBP2 |
| 6 | CACNA1D | 56 | SNAP25 | 106 | SYT10 |
| 7 | CACNA1F | 57 | SNAP29 | 107 | SYT15 |
| 8 | AP1G1 | 58 | STX12 | 108 | SYT4 |
| 9 | AP1M1 | 59 | STX18 | 109 | SYT5 |
| 10 | AP1M2 | 60 | STX4 | 110 | SYT9 |
| 11 | AP1S1 | 61 | STX8 | 111 | VAMP1 |
| 12 | CACNA1B | 62 | STXBP5 | 112 | VAMP3 |
| 13 | CACNA1C | 63 | STXBP6 | 113 | VAMP4 |
| 14 | CAV2 | 64 | SYT1 | 114 | VAMP8 |
| 15 | DNM1 | 65 | SYT13 | 115 | VPS13B |
| 16 | EHD1 | 66 | SYT16 | 116 | VPS33B |
| 17 | EHD4 | 67 | SYT17 | 117 | ESYT2 |
| 18 | ESYT3 | 68 | SYT2 | 118 | STXBP3 |
| 19 | HPS4 | 69 | SYT3 | 119 | STXBP4 |
| 20 | HUWE1 | 70 | SYT6 | 120 | AP1AR |
| 21 | PLIN3 | 71 | SYT7 | 121 | AP1S2 |
| 22 | PREB | 72 | SYT8 | 122 | AP3S2 |
| 23 | STX11 | 73 | VAPB | 123 | AP4S1 |
| 24 | STX1A | 74 | AP1S3 | 124 | ARHGAP32 |
| 25 | STX1B | 75 | AP2A2 | 125 | CACNA1H |
| 26 | SYNRG | 76 | AP3B2 | 126 | CDKN2A |
| 27 | SYT11 | 77 | AP3M1 | 127 | CLTC |
| 28 | AP1B1 | 78 | CACNA1E | 128 | COPB1 |
| 29 | AP2B1 | 79 | CACNA1I | 129 | COPG2 |
| 30 | AP2S1 | 80 | CACNA1S | 130 | COPZ2 |
| 31 | AP3B1 | 81 | CAV1 | 131 | HPS3 |
| 32 | AP3M2 | 82 | COPE | 132 | HPS6 |
| 33 | AP3S1 | 83 | COPZ1 | 133 | LYST |
| 34 | AP4B1 | 84 | EHD3 | 134 | SAR1A |
| 35 | AP4E1 | 85 | HPS1 | 135 | SEC24B |
| 36 | AP4M1 | 86 | ITGA3 | 136 | STX17 |
| 37 | ARF1 | 87 | LMAN1 | 137 | SYT12 |
| 38 | ARHGAP10 | 88 | PTGES2 | 138 | SYT14 |
| 39 | CACNA1A | 89 | RAB3A | 139 | TBC1D15 |
| 40 | CACNA1G | 90 | RACGAP1 | 140 | VAMP2 |
| 41 | CAV3 | 91 | RASA1 | 141 | VAMP7 |
| 42 | CLTA | 92 | RGS6 |  |  |
| 43 | CLTB | 93 | SAR1B |  |  |
| 44 | COPA | 94 | SEC23A |  |  |
| 45 | COPB2 | 95 | SEC31A |  |  |
| 46 | COPG1 | 96 | SNAP23 |  |  |
| 47 | DNM2 | 97 | STX10 |  |  |
| 48 | DNM3 | 98 | STX19 |  |  |
| 49 | DTNBP1 | 99 | STX2 |  |  |
| 50 | EHD2 | 100 | STX3 |  |  |

**Supplementary table 2.**
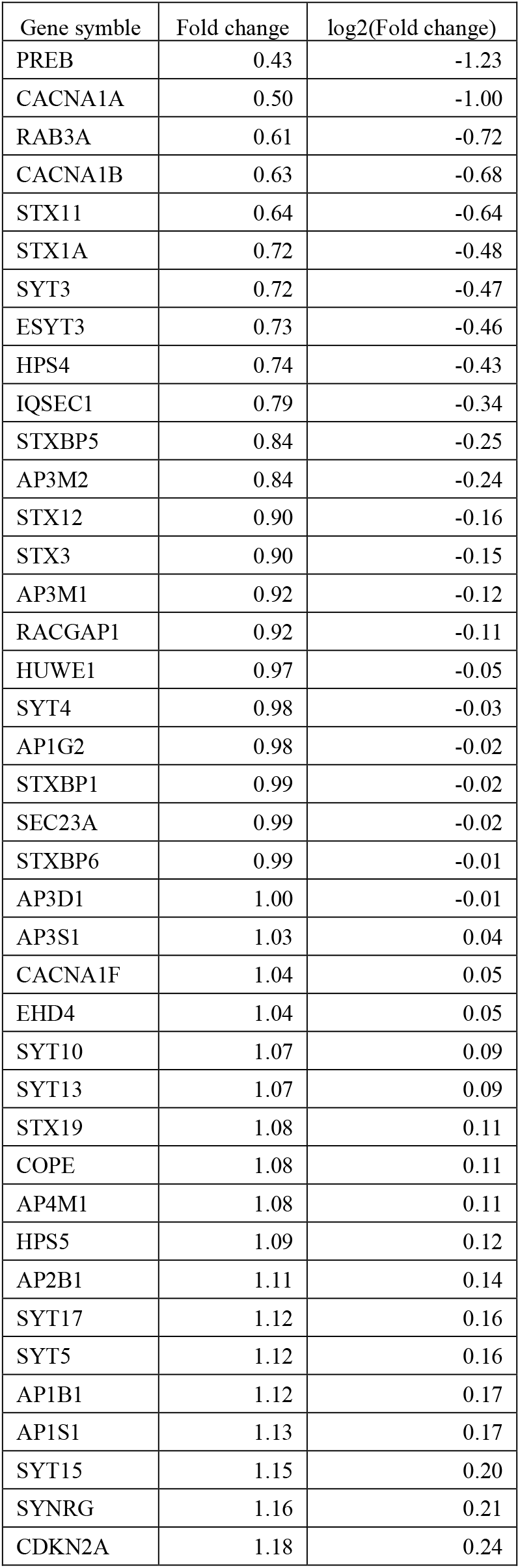
40membrane trafficking genes, knocking out which relieved CTSG induced upregulation of cleaved PARP to a level stronger than knocking out RAGE.

**Supplementary table 3.** Top 10 increased and decreased genes in neutrophil conditional medium treated DLD1 cells compared to fresh medium cultured DLD1 cells.

| Gene | sgRNA | CM-N\|beta | CM-N\|z | CM-N\|p-value |
| --- | --- | --- | --- | --- |
| PSMB7 | 4 | 0.68592 | 0.90644 | 0 |
| TMEM88B | 4 | 0.5362 | 1.0924 | 5.23E-05 |
| NDC80 | 4 | 0.50108 | 0.79144 | 0.00010463 |
| DEFB106B | 2 | 0.50077 | 0.76842 | 0.0044468 |
| RSRC1 | 4 | 0.49446 | 1.0511 | 0.00015694 |
| NRF1 | 4 | 0.49007 | 1.1138 | 0.00015694 |
| SCD | 4 | 0.48661 | 0.88939 | 0.00015694 |
| RBM22 | 4 | 0.47152 | 0.69873 | 0.00015694 |
| CDK1 | 4 | 0.46026 | 0.87898 | 0.00015694 |
| ISY1-RAB43 | 4 | 0.44773 | 1.1895 | 0.00015694 |
| KDELR2 | 4 | -0.43261 | -1.1169 | 0.0003662 |
| INCENP | 4 | -0.43795 | -1.2528 | 0.0003662 |
| POLR2A | 4 | -0.44258 | -1.2716 | 0.0003662 |
| SETD8 | 4 | -0.45891 | -1.6657 | 0.00026157 |
| DROSHA | 4 | -0.46408 | -1.9457 | 0.00026157 |
| RPS15 | 4 | -0.48098 | -1.7574 | 0.00026157 |
| SNRPD3 | 4 | -0.51662 | -1.6884 | 0.00026157 |
| WASH1 | 4 | -0.51714 | -2.0189 | 0.00026157 |
| SNRPA1 | 4 | -0.5493 | -1.424 | 0.00020926 |
| ZNF830 | 4 | -0.61166 | -2.0198 | 0.00010463 |

